# *Foxf1* is required for the specification and maintenance of pulmonary capillary identity

**DOI:** 10.64898/2026.06.16.732674

**Authors:** Yun Liu, Amanda Ceas, David C. Arteaga, Jonathan D. Bywaters, Tanya V. Kalin, Vladimir V. Kalinichenko, Luisa Morales-Nebreda, Lisandra Vila Ellis

**Affiliations:** Department of Cell & Developmental Biology, Northwestern University Feinberg School of Medicine, Chicago, IL, USA; UT Health Science Center, San Antonio, Texas, USA; Department of Child Health, Phoenix Children’s Research Institute, University of Arizona College of Medicine-Phoenix, Phoenix, AZ, USA; Phoenix Children’s Center for Cancer and Blood Disorders, Phoenix Children’s Hospital, Phoenix AZ, USA; Division of Neonatology, Phoenix Children’s Hospital, University of Arizona College of Medicine – Phoenix, Phoenix, AZ, USA; Division of Pulmonary and Critical Care Medicine, Department of Medicine, Northwestern University Feinberg School of Medicine, Chicago, IL, USA; Simpson Querrey Lung Institute for Translational Science, Northwestern University Feinberg School of Medicine, Chicago, IL, USA

**Author notes:** corresponding author **CONTACT:** Lisandra Vila Ellis, 303 East Superior Street, Simpson Querrey 8-301, Chicago, Illinois 60611.

**Keywords:** Foxf1, lung vascular development, lung endothelial cells

## Abstract

The pulmonary microvasculature consists of two transcriptionally distinct capillary (CAP) endothelial populations, CAP1 and CAP2 cells, that form the alveolar capillary network. While CAP1 and CAP2 are functionally distinct, the transcriptional mechanisms that specify and maintain these endothelial fates remain poorly understood. The Forkhead transcription factor *Foxf1* regulates vascular development and is implicated in the neonatal disease alveolar capillary dysplasia with misalignment of pulmonary veins (ACDMPV). However, its role in the specification of pulmonary endothelial subtypes has not been defined. Using single cell ATAC-sequencing and RNA-sequencing of developing mouse lung endothelial cells (ECs), we found that *Foxf1* is broadly expressed across pulmonary ECs but exhibits preferential chromatin accessibility in CAP2 ECs. Endothelial deletion of *Foxf1* impaired CAP2 specification and disrupted CAP1 identity, unexpectedly leading to the emergence of a mutant CAP1-like population with a macrovascular transcriptional signature. Adult endothelial deletion of *Foxf1* similarly resulted in loss of capillary identity, demonstrating that *Foxf1* is required for capillary fate maintenance. Mutant CAP1 ECs upregulated matrix remodeling genes and exhibited altered communication with neighboring mesenchymal populations, suggesting a potential role in disease development. Consistent with these findings, this mutant CAP1 transcriptional signature was present in human lungs with ACDMPV. This work identifies *Foxf1* as a key transcriptional regulator for the specification and active maintenance of pulmonary capillary EC fate across the lifespan, and positions capillary identity loss as a driver of vascular remodeling.

## INTRODUCTION

Pulmonary capillaries, once considered a homogenous population, are now recognized for their heterogeneity. The alveolar vascular network consists of primarily two alveolar capillary (CAP) endothelial subtypes, CAP1 and CAP2 cells, which differ in their transcriptome, morphology, localization and potentially function.^1,2^ These endothelial cells (ECs) arise from a common progenitor, and during the saccular stage of lung development, epithelial-derived Vascular endothelial growth factor A (VEGFA) promotes the specification of CAP2 cells.^1^ CAP2 cells seem poised for gas exchange due to their proximity and interaction with alveolar type 1 (AT1) cells, as well as their gene expression, while CAP1 cells can serve as progenitors during neonatal injury and give rise to CAP2 cells.^1–3^

Transcription factors are key regulators of cell fate in many developmental contexts. For instance, NK2 Homeobox 1 (NKX2-1) is essential for lung development, acting as a pioneer factor that regulates alveolar epithelial specification by recruiting distinct co-factors in a cell type-specific manner.^4,5^ In the vasculature, venous versus arterial specification is better understood than capillary fates; however, capillaries are more plastic and adapt to their vascular bed, serving as the main source of endothelial heterogeneity.^6–8^ In the lungs, the transcriptional mechanisms that establish distinct endothelial fates in the lung remain unknown.

The Forkhead family of transcription factors is known to regulate tissue morphogenesis and disease states.^9^ Among this family, *Forkhead Box F1 (Foxf1)* is required for mesoderm differentiation and vasculogenesis, and plays a critical role in foregut and lung development.^10,11^ Complete loss of *Foxf1* results in embryonic lethality around embryonic (E) day 9, while overexpression leads to lung hypoplasia and vascular defects.^11,12^ Importantly, mutations in human FOXF1 cause alveolar capillary dysplasia with misalignment of pulmonary veins (ACDMPV), a typically fatal neonatal lung disease.^13^ Affected infants present with severe hypoxemia and pulmonary hypertension (PH) at birth, although rare, atypical cases have been described in which initially asymptomatic neonates later develop PH.^13–16^ Because ACDMPV is caused by heterozygous FOXF1 mutations, previous studies have largely focused on haploinsufficiency and disease-associated point mutations.^17–19^ However, the role of FOXF1 in the specification and maintenance of pulmonary capillary endothelial identity remains unknown.

## RESULTS

### FOXF1 emerges as a candidate regulator of pulmonary capillary endothelial fate

To identify potential regulators of capillary fate in an unbiased manner, we analyzed single cell ATAC-sequencing data (scATAC-seq) of postnatal (P) mouse lung ECs (**Fig. 1A**).^20^ By surveying the chromatin accessibility of canonical marker genes, we identified endothelial subtypes including CAP1, CAP2 and macrovascular ECs (**Fig. 1B**). Motif enrichment analysis revealed the *Fox* family as the top candidate for CAP2 specification, leading us to focus on FOXF1 due to its established role in lung vascular development.^18,21^ To define the cell type-specific expression of *Foxf1*, we analyzed published single cell RNA-sequencing (scRNA-seq) datasets spanning embryonic, early postnatal and adult mouse lungs and identified the major cell lineages based on established markers (**Fig. 1D, Fig. S1A**).^1^ Although endothelial *Foxf1* has been previously noted in a subset of KIT+ ECs,^18,22^ our scRNA-seq analysis revealed broad *Foxf1* expression in all ECs, with higher levels in CAP2 cells during perinatal stages (**Fig. 1E, Fig. S1B-D**). Outside the endothelium, *Foxf1* expression was largely restricted to myofibroblasts and pericytes as previously described (**Fig. 1E, Fig. S1E-G**).^18^ To validate this expression pattern, we immunostained lung sections at various timepoints using ETS Transcription Factor ERG (ERG) as a pan-endothelial marker, and Platelet Derived Growth Factor Receptor Alpha or Beta (PDGFRA or PDGFRB) for myofibroblasts and pericytes, respectively (**Fig. 1F-G, Fig. S1H, I**). Immunostaining confirmed FOXF1 expression in both endothelial and mesenchymal compartments across developmental stages. While endothelial FOXF1 expression persisted into adulthood, non-endothelial FOXF1 expression was markedly reduced, consistent with the loss of alveolar myofibroblasts in the adult lung (**Fig. 1F**).^23^ Together, these findings identified FOXF1 as a candidate regulator of pulmonary capillary endothelial fate and suggested that, despite preferential accessibility in CAP2 ECs, FOXF1 may function more broadly across endothelial subtypes.

**Figure 1.**
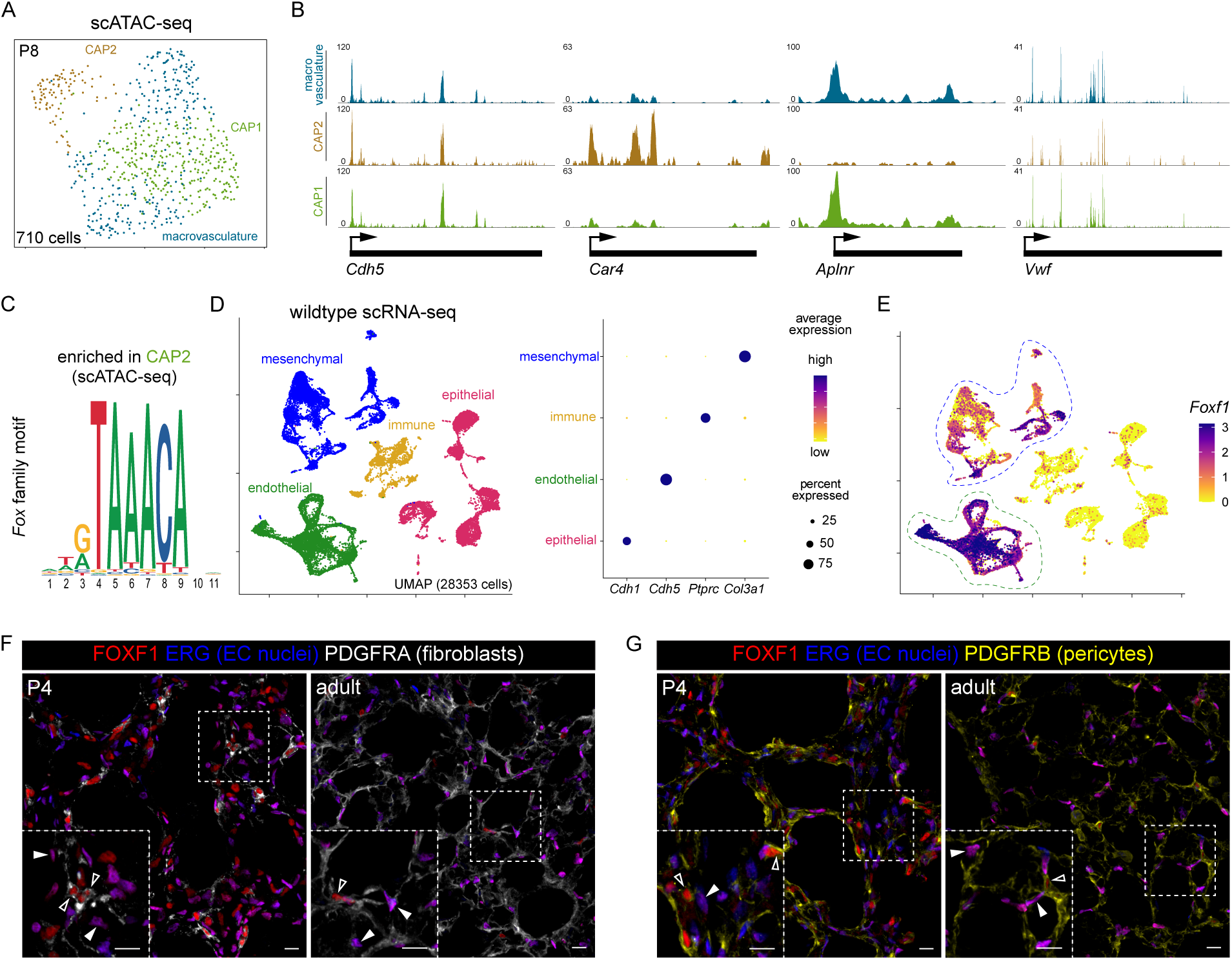
*Foxf1* is enriched in pulmonary ECs and highly accessible in CAP2 ECs. (**A**) scATAC-seq UMAP showing mouse lung EC populations at P8, color-coded by EC population. (**B**) Coverage plots of scATAC-seq data showing differential accessibility of marker genes *Cdh5* (pan-endothelial), *Car4* (CAP2), *Aplnr* (CAP1), and *Vwf* (macrovasculature) allowing cell-type identification. (**C**) Motif sequence enriched in CAP2 ECs revealing *Fox* family members as a top candidate. (**D**) scRNA-seq UMAP of lungs from wildtype mice colored by lineage (left). Dot plot showing markers used to identify each lineage (right). (**E**) Feature plot showing *Foxf1* expression in the lung. Dashed lines highlight endothelial (green) and mesenchymal (blue) expression of *Foxf1*. (**F-G**) Section immunostaining of wildtype P4 and adult lungs validates expression of FOXF1 in fibroblasts (PDGFRA-wrapped FOXF1+ nuclei, white open arrowhead), ECs (FOXF1+ERG+ nuclei, white filled arrowhead) or pericytes (PDGFRB-wrapped FOXF1+ nuclei, white open arrowhead). Boxed regions are magnified. P, postnatal day. Scale bars = 10 μm.

### Loss of *Foxf1* disrupts CAP1 and CAP2 fate and promotes macrovascular identity

To functionally test the role of *Foxf1* in the specification of EC subtypes, we deleted *Foxf1* at birth using the inducible pan-endothelial driver *Cdh5-CreER* (hereafter referred to as *Foxf1^EC^*).^24,25^ Importantly, neonatal *Foxf1^EC^*mice died within 5–7 days post-deletion, precluding functional assessments and highlighting the essential role of FOXF1 during early postnatal life. Thus, we injected newborn pups at birth and harvested 4 days after injection at P4 (**Fig. 2A**). scRNA-seq of sorted lung cells identified all major pulmonary cell populations and confirmed selective loss of endothelial, but not mesenchymal, *Foxf1* expression (**Fig. S2A–E**). We then subclustered the endothelial populations and observed a striking transcriptional reprogramming of the *Foxf1^EC^* endothelium (**Fig. 2B**). Using known markers, we identified each EC cluster (**Fig. 2C**). In line with the enrichment of FOXF1 accessibility in CAP2 cells, CAP2 specification was impaired, resulting in failure of this population to segregate in the *Foxf1^EC^* mutant. The population of early CAP2 ECs also increased in the *Foxf1^EC^* mutant suggesting blunted differentiation of CAP2 cells. Unexpectedly, CAP1 cells were nearly absent in the mutant lungs and replaced by a CAP1-like population (**Fig. 2B, D**). Notably, EC proliferation seemed unchanged (**Fig. 2D, Fig. S3A**). Differential gene expression analysis between all ECs in the control and *Foxf1^EC^*lungs showed significant downregulation of canonical marker genes for CAP2 cells such as *carbonic anhydrase 4 (Car4)* and *Fin Bud Initiation Factor Homolog (Fibin)*, and CAP1 genes like *Wnt Inhibitory Factor 1 (Wif1)*,^1^ suggesting overall loss of capillary identity (**Fig. 2E, Dataset S1**).

**Figure 2.**
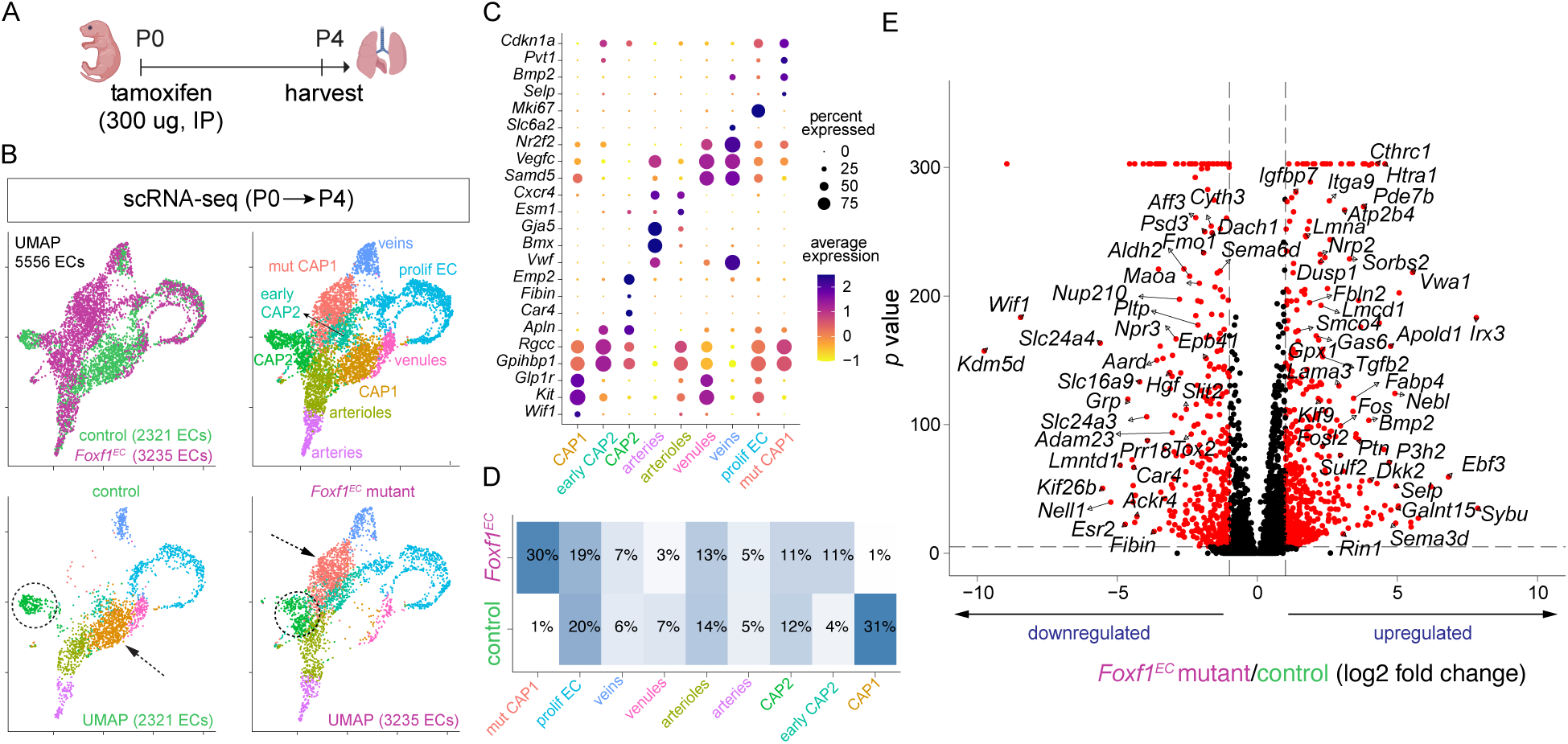
scRNA-seq shows transcriptional shift in *Foxf1* mutant endothelium reflecting capillary loss. (**A**) Schematic of P4 experiment created with *BioRender*. (**B**) UMAPs of sorted lung ECs from P4 control and *Foxf1^EC^* mutant mice colored by condition (top left), EC subpopulation (top right) and split by condition (bottom). Dashed circles highlight CAP2 shift and dashed arrows highlight loss of CAP1 cluster and emergence of mutant-specific CAP1 population. (**C**) Dot plot showing markers used to identify each population. (**D**) Percentages of EC populations between control and *Foxf1^EC^* lungs. (**E**) Volcano plot showing differentially expressed genes between mutant and control ECs (**Dataset S1**). IP, intraperitoneal; P, postnatal day; mut, mutant; prolif, proliferative.

To distinguish between a role in developmental specification vs maintenance of established endothelial identities, we induced *Foxf1* deletion in adult mice and compared the resulting vascular phenotype with that observed following neonatal deletion (**Fig. 3A, 2A**). Adult and neonatal ECs express FOXF1 at similar rates, and we achieved efficient deletion at both stages (**Fig. 3B**). Total EC number remained comparable between control and *Foxf1^EC^* mutant (**Fig. 3B**) and cleaved-caspase 3 (CASP3) staining reveal no changes in endothelial apoptosis (**Fig. S3B**). These findings indicate that loss of *Foxf1* does not result in EC loss, suggesting that the vascular changes observed may arise from altered endothelial state rather than impaired survival. Total vascular surface area was significantly reduced at both timepoints (**Fig. 3C**), indicating that although EC number remained unchanged, endothelial morphology could be altered, resulting in a less complex vascular network. Given the large size and web-like morphology of CAP2 ECs,^1^ we hypothesized that reduced CAP2 development contributed to the decreased vascular surface area. To test this, we stained for the CAP1 and CAP2 markers Plasmalemma Vesicle Associated Protein (PLVAP) and CAR4, respectively. Both markers were significantly reduced in neonatal and adult *Foxf1^EC^* mutants, as quantified by CAP1 and CAP2 surface area (**Fig. 3D**). Because lung development proceeds in a proximal-to-distal manner, newly specified CAP2 cells are expected to localize near the distal lung edge. We therefore measured the distance between CAP2 cells and the lung periphery as a surrogate for ongoing CAP2 specification. This distance was significantly increased in *Foxf1^EC^* mutants, consistent with impaired distal capillary development (**Fig. S3C**). Although CAR4 staining was markedly reduced following neonatal deletion, residual CAR4+ ECs remained, likely representing CAP2 cells that had been specified prior to *Foxf1* deletion at birth. To directly test whether FOXF1 is required for CAP2 specification, we deleted *Foxf1* at E15, before CAP2 specification occurs around E18 (**Fig. 3E**).^1^ These lungs showed near-complete absence of CAR4 expression, with only rare escaper cells remaining (**Fig. 3F**). Finally, we generated module scores for each EC population using their top 50 maker genes in control conditions. Capillary EC scores revealed loss of canonical genes specific to CAP1 and CAP2 in the *Foxf1^EC^* mutant (**Fig. 3G, Dataset S2**).

**Figure 3.**
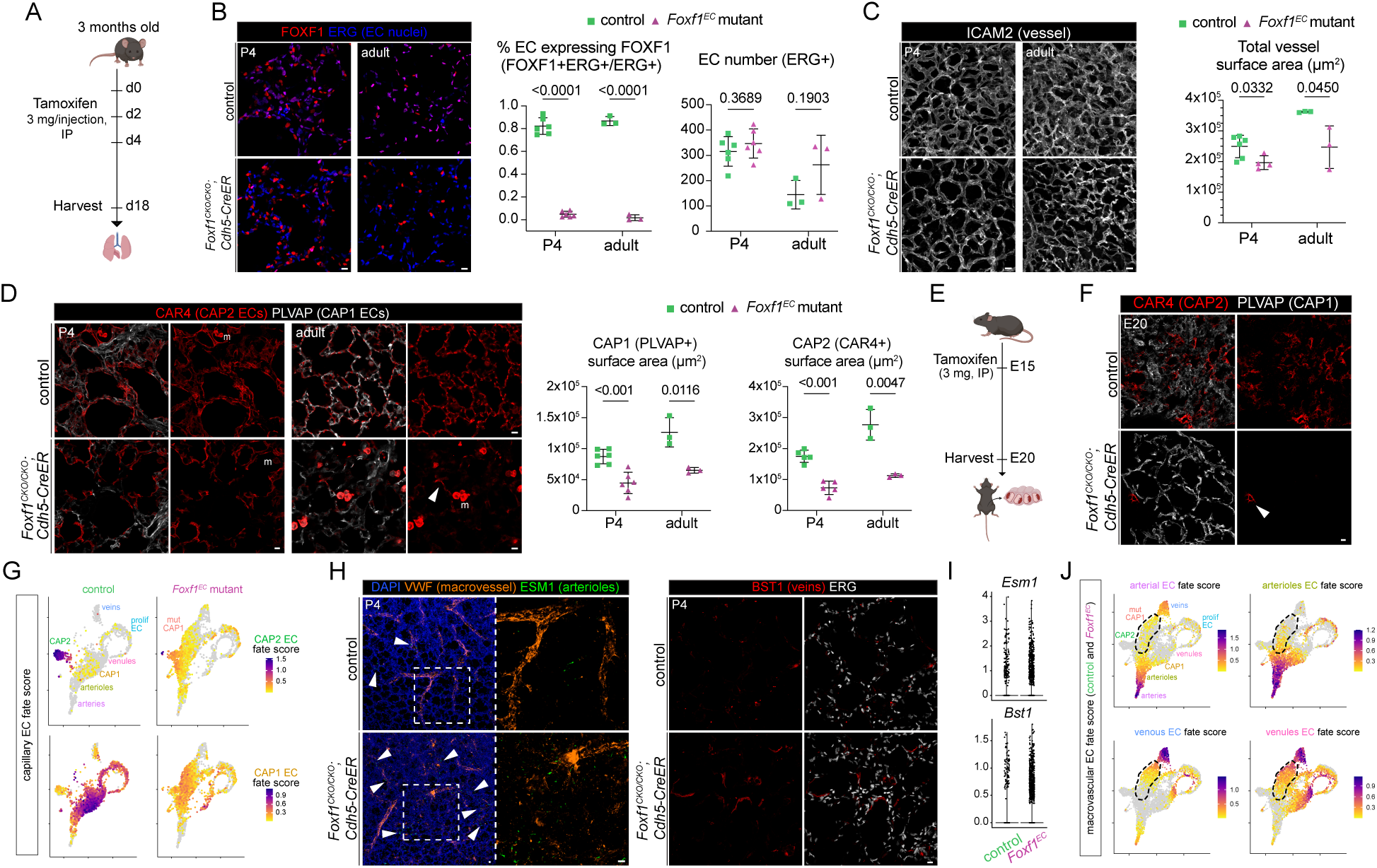
Endothelial deletion of *Foxf1* leads to loss of capillary identity and upregulation of macrovascular EC fate. (**A**) Schematic of adult experiment. **(B**) Section immunostaining of P4 and adult lungs validates loss of FOXF1 in ECs (ERG+ nuclei). Quantification shows efficient deletion of FOXF1 and no significant change in EC number. (**C**) Wholemount immunostaining of P4 and adult lungs showing reduction of ICAM2 in *Foxf1^EC^* mutants along with quantification of total vessel surface area using ICAM2. (**D**) Section immunostaining of P4 and adult lungs showing decreased PLVAP (CAP1) and CAR4 (CAP2) expression in mutants. Adult *Foxf1^EC^* lungs have rare CAR4+ endothelial expression (arrowhead). Quantification of CAP1 and CAP2 vessel surface area reveals significant loss of both capillary populations in P4 and adult mutant. (**E**) Schematic of embryonic experiment. (**F**) Section immunostaining of E20 lung showing simplification of PLVAP+ vascular network (CAP1) and loss of CAR4 (CAP2) except for very few and rare escapers (arrowhead). (**G**) Feature plots of capillary module scores reveal loss of capillary signature by CAP1 and CAP2 ECs in *Foxf1^EC^*. (**H**) Immunostaining of P4 lungs reveals ectopic expression of macrovascular markers VWF, ESM1, and BST1 in distal alveolar regions of the mutant. (**I**) Violin plots showing corresponding transcriptional increase of *Esm1* and *Bst1* in *Foxf1^EC^*mutant. (**J**) Feature plots of macrovascular module scores reveal gain of macrovascular gene signatures, particularly venous/venular, by mutant CAP1 cells (dashed). Each quantification datapoint represents the average of 3 distinct images within 1 mouse. All *p*-values were calculated using Student’s t test. Boxed regions are magnified. IP, intraperitoneal; m, macrophage; P, postnatal day; E, embryonic day; mut, mutant; prolif, proliferative. Schematics were created with *BioRender.* Scale bars = 10 μm.

To determine whether loss of capillary identity was accompanied by acquisition of alternative vascular fates, we examined expression of macrovascular markers Von Willebrand Factor (VWF, pan-macrovascular), Endothelial Cell Specific Molecule 1 (ESM1, lung arterioles), and Bone Marrow Stromal Cell Antigen 1 (BST1, veins) (**Fig. 3H**). These markers were found ectopically expressed in distal alveolar regions, and both *Esm1* and *Bst1* were upregulated in mutant lungs (**Fig. 3H, I**). Indeed, macrovascular fate module scores were selectively enriched in mutant CAP1 cells, with prominent enrichment of venous and venular signatures (**Fig. 3J, Dataset S2**). These findings demonstrate that FOXF1 is required for the specification and maintenance of pulmonary capillary endothelial identity while repressing macrovascular gene programs. More broadly, they reveal that pulmonary capillary fate is not fixed after specification but instead requires continuous FOXF1-dependent maintenance.

### Mutant CAP1 cells acquire a vascular remodeling program and exhibit altered mesenchymal communication

An unexpected consequence of endothelial *Foxf1* deletion was the emergence of a mutant CAP1 population that had largely lost canonical CAP1 transcriptional identity. To investigate the origin and fate of these cells, we performed trajectory analysis which suggested that mutant CAP1 cells arise from CAP1 or early CAP2 cells, and some acquire a venous fate while others remain in this mutant state (**Fig. 4A**). Further supporting a venous-like signature, the top marker genes enriched in mutant CAP1 cells showed shared expression with venous ECs (**Fig. 4B**). One of the top upregulated genes in mutant CAP1 cells was *Cyclin Dependent Kinase Inhibitor 1A (Cdkn1a* or *p21)*. Although only ∼50% of mutant CAP1 cells express P21, this marker enabled *in situ* validation of the mutant population (**Fig. 4C**). Due to the aberrant transcriptional state of these cells, we asked whether they were undergoing senescence. To address this, we examined expression of senescence-associated secretory phenotype (SASP) genes and calculated a senescence score using the Mayo senescence (senMayo) gene set (**Fig. S4A**).^26^ Neither SASP nor senMayo scores were significantly elevated in mutant CAP1 cells. In addition, P21+ ECs retained Lamin B1 (LMNB1) expression, a hallmark protein lost during senescence, arguing against a canonical senescent phenotype (**Fig. S4B**).^27^ This is consistent with our previous findings that Tumor Protein P53 (p53) target genes, including *Cdkn1a*, are induced during endothelial fate transitions to maintain lineage fidelity.^3^ Interestingly, other significantly upregulated genes including *Bone Morphogenetic Protein 2 (Bmp2)*, *Transforming Growth Factor Beta Receptor 2 (Tgfbr2)* and *EBF Transcription Factor 1 (Ebf1)*, have been described in association with PH, a common clinical feature of ACDMPV (**Fig. 4D**).^13,14,28,29^ Gene ontology (GO) analysis of mutant CAP1 marker genes revealed enrichment of pathways involved in extracellular matrix organization and fibroblast-regulatory programs, suggesting a role for these ECs in vascular remodeling (**Fig. 4E**).

**Figure 4.**
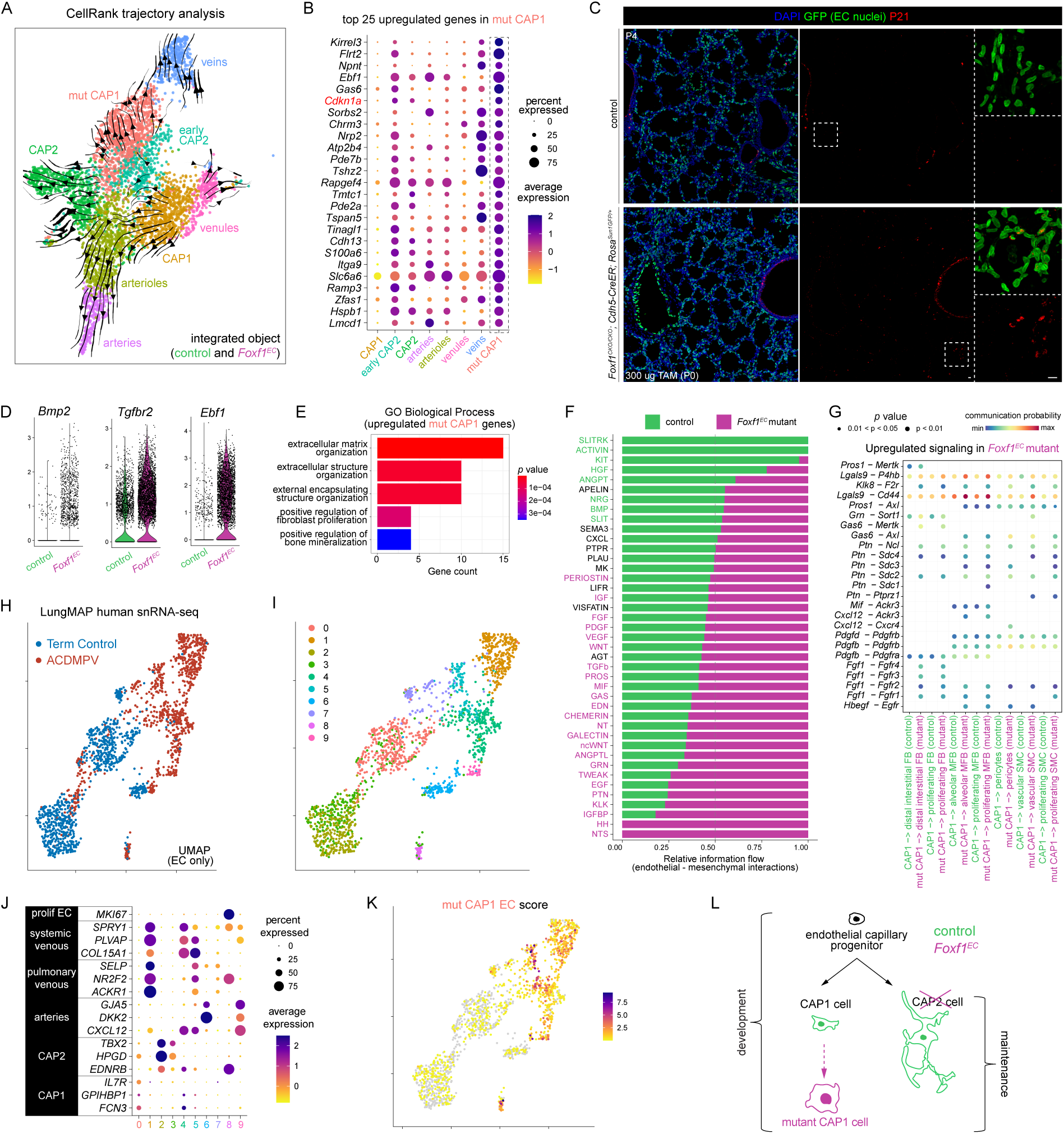
Mutant CAP1 ECs undergo vascular remodeling and may play a role in disease. (**A**) CellRank trajectory analysis suggests that CAP1 and early CAP2 ECs transition to mutant CAP1 cells, which remain in this state or transition to venous ECs. (**B**) Dot plot of the top 25 upregulated genes in mutant CAP1 ECs, including *Cdkn1a* (*p21*). (**C**) Section immunostaining of P4 lung sections showing increased P21 expression in *Foxf1* mutant ECs. *Rosa^SUN1GFP^* labels ECs and can be co-localized with P21 indicating mutant CAP1 cells. (**D**) Violin plots exhibit upregulation of PH-associated genes *Bmp2*, *Tgfbr2*, and *Ebf1* in *Foxf1^EC^* mutant. (**E**) GO Biological Process analysis of genes significantly upregulated in mutant CAP1 ECs shows remodeling-associated programs. (**F**) CellChat analysis showing changes in endothelial and mesenchymal signaling in *Foxf1^EC^* mutant. (**G**) Dot plot of predicted, upregulated ligand-receptor interactions in *Foxf1^EC^* mutant with CAP1 as sender and mesenchymal subpopulations as targets. (**H**) UMAP of human lung ACDMPV snRNA-seq data colored by condition or (**I**) by Seurat cluster. (**J**) Dot plot of marker genes used to identify each endothelial subpopulation in human snRNA-seq data. (**K**) Feature plot of mutant CAP1 score shows mutant CAP1–like transcriptional profile in ACDMPV ECs. (**L**) Schematic of proposed model showing normal capillary differentiation requires FOXF1 and the disruption of CAP1 and CAP2 differentiation without FOXF1 in developmental and adult lungs. Boxed regions are magnified. TAM, tamoxifen; P, postnatal day; mut, mutant; prolif, proliferative; FB, fibroblasts; MFB, myofibroblasts; SMC, smooth muscle cells. Scale bars = 10 μm.

Because many of the enriched pathways were associated with extracellular signaling and tissue remodeling, we next investigated whether mutant CAP1 cells exhibit altered communication with neighboring mesenchymal populations (**Fig. 4F**). Mutant CAP1 cells upregulated several genes implicated in extracellular matrix remodeling, including *Von Willebrand Factor A Domain Containing 1 (Vwa1)* and components of the TGF-β signaling pathway (**Fig. 2E**),^30^ suggesting acquisition of a signaling program capable of influencing the surrounding alveolar niche. Analysis of ligand-receptor interactions revealed enhanced communication between mutant CAP1 cells and multiple mesenchymal populations, including fibroblasts, myofibroblasts, pericytes, vascular smooth muscle cells, and proliferating mesenchymal cells (**Fig. 4F**). Further, the outgoing signals from mutant CAP1 cells showed increased activity of pathways involved in stromal regulation and tissue remodeling, including TGF-β, PDGF, Fibroblast Growth Factor (FGF), and Pleiotrophin (PTN) signaling (**Fig. 4G, Dataset S3**). In contrast, signaling pathways reduced in mutant CAP1 cells included semaphorin and BMP signaling which have previously been identified as downstream targets of FOXF1 (**Fig. S4C, Dataset S4**).^18,31,32^ This reprogramming suggests that capillary endothelial fate maintenance is necessary to preserve normal endothelial-mesenchymal communication within the alveolar niche and that loss of endothelial identity promotes remodeling-associated signaling programs.

### Mutant CAP1 transcriptional state is conserved in human ACDMPV lungs

Finally, we analyzed lung ECs from published single nuclei RNA-sequencing (snRNA-seq) data of healthy donors and patients with ACDMPV (**Fig. 4H**).^32^ Among these ECs, we detected 9 clusters that encompass the main endothelial subtypes identified by canonical marker genes (**Fig. 4I, J**). Similar to our mouse model, ACDMPV lung exhibited a loss of normal capillary endothelial populations (**Fig. 4H, I**). To determine whether the mutant CAP1 state was conserved across species, we generated a module score based on the mutant CAP1 transcriptional signature identified in *Foxf1^EC^*mice. This signature was significantly enriched in ECs from ACDMPV lungs (**Fig. 4K, Dataset S2**), indicating that the aberrant capillary state observed following *Foxf1* endothelial loss is also present in human disease.

In conclusion, these data demonstrate that FOXF1 is required to establish and maintain lung capillary EC identity (**Fig. 4L**). These findings further indicate that FOXF1 plays cell type-specific roles within the pulmonary endothelium, suppressing macrovascular fate in the capillaries. Furthermore, we show that a novel CAP1 dysfunctional state arises in the absence of *Foxf1* characterized by altered endothelial-mesenchymal communication and remodeling-associated signaling. The presence of this transcriptional state in human ACDMPV lungs highlights its potential contribution to disease pathogenesis.

## DISCUSSION

Pulmonary capillaries are composed of transcriptionally and functionally distinct endothelial populations, yet the mechanisms that establish and maintain this heterogeneity have remained poorly understood. Here, we identify FOXF1 as a critical regulator of lung capillary EC identity, required for both specification during development and maintenance of capillary identity in the adult lung. These findings extend previous work implicating FOXF1 in lung vascular development by demonstrating that its loss produces distinct transcriptional consequences across capillary endothelial subtypes, resulting in loss of capillary identity and acquisition of macrovascular programs.

A key finding of this study is the emergence of a distinct CAP1-like population following *Foxf1* endothelial deletion, characterized by venous and macrovascular gene expression, extracellular matrix remodeling signatures, and upregulation of stress-associated genes, including P21. The acquisition of these macrovascular features suggests that capillary ECs retain latent plasticity toward alternative vascular identities, indicating that capillary endothelial identity is not terminally fixed but requires continuous transcriptional regulation. Moreover, these findings show that FOXF1 stabilizes capillary-specific transcriptional programs and restrict activation of alternative endothelial fates. Conceptually, this model parallels lineage control mechanisms described in other lung compartments; for example, NKX2-1 directs alveolar epithelial cell fate through cell type-dependent chromatin binding.^4,5^

Importantly, our findings have direct relevance to pulmonary vascular disease. Mutations in FOXF1 are causative in ACDMPV, a lethal neonatal disorder characterized by severe vascular abnormalities.^13,32,33^ The mutant CAP1-like population that emerges following *Foxf1* deletion is enriched for pulmonary hypertension-associated genes, exhibits altered endothelial-mesenchymal communication, and upregulates vascular remodeling programs. These findings demonstrate that endothelial loss of *Foxf1* alone is sufficient to induce remodeling-associated changes, supporting the idea that disruption of endothelial fate may act as a driver of tissue remodeling rather than simply a consequence of disease progression. The identification of a mutant CAP1-like endothelial state in both the mouse model and human ACDMPV lungs suggests that disease pathogenesis may involve not only impaired capillary development but also the emergence of aberrant endothelial identities. This shift in perspective highlights endothelial misspecification as a potential driver of vascular remodeling and PH. Specifically, a transcriptional shift towards venous fate could provide an explanation to the unknown origin of the misalignment of pulmonary veins observed in this disease.

While these findings provide new insights for understanding capillary endothelial identity, several limitations should be considered. The mutant CAP1 population is defined primarily by single cell transcriptomic profiling, although its presence is supported by *in situ* validation and consistent transcriptional signatures across datasets. In addition, early lethality following neonatal *Foxf1* deletion limits functional assessment at this stage. Although adult deletion demonstrates a requirement for maintenance of capillary identity, mice do not exhibit rapid lethality within the time frame examined (two weeks post-deletion), suggesting distinct physiological consequences between neonatal and adult loss of *Foxf1*. Finally, the direct molecular mechanisms by which FOXF1 regulates capillary endothelial programs, including its potential cell type-specific co-factors, remain to be defined.

In summary, our data identify FOXF1 as a central regulator of pulmonary capillary EC fate. Loss of *Foxf1* leads to disruption of capillary programs and the emergence of a dysfunctional endothelial state associated with disease. These findings provide a novel conceptual framework for understanding endothelial heterogeneity and its dysregulation in pulmonary vascular disease.

## MATERIALS AND METHODS

### Antibodies

See supplementary information (**Dataset S5**).

### Experimental mouse models

The following experimental mice were used: *Foxf1^fl/fl^* (Dr. Vladimir Kalinichenko and Dr. Tanya V. Kalin), *Cdh5-CreER* (Dr. Ralf Adams, CancerTools UK), *Rosa^Sun1GFP^* (JAX 021039). ^24,25,34^ For timed pregnancies, the day of observing a vaginal plug was designated as E0.5. Tamoxifen (Sigma, T5648) was dissolved in corn oil (Sigma, C8267) at a 10 mg/mL concentration, and administered to pups via intraperitoneal injection to induce Cre-Lox recombination when required. Time and dose of injection is indicated in figures. Mice of both sexes were used and the number of samples used for quantification is indicated in figure legends. All protocols used for this research complied with and were approved by the Institutional Animal Care and Use Committee regulations of Northwestern University.

### Lung harvest

Post-natal lungs were harvested as previously described.^1,3^ Briefly, mice were anesthetized with avertin (T48402, Sigma) and perfused with phosphate-buffered saline (PBS, Thermo Scientific, BP6651) via the right ventricle. Then, the trachea was cannulated and the lungs were inflated with 0.5% paraformaldehyde (PFA, Thermo Scientific, J19943.K2) in PBS at 25 cm H_2_O pressure. Pregnant female mice were euthanized, the uterus was dissected and kept in cold PBS. Embryos were dissected from the uterus while submerged in PBS using a dissection scope (Olympus SZ51 Stereomicroscope), then embryonic lungs were harvested. Both post-natal and embryonic lungs were fixed in 0.5% PFA for 3-6 hours at room temperature and washed in PBS at 4°C overnight.

### Section immunostaining

Section immunostaining was performed as previously described.^1,3^ Fixed lung lobes were cryoprotected in PBS in 20% sucrose with 10% optimal cutting temperature compound (OCT, Sakura Finetek, 4583) at 4°C overnight and then frozen in OCT blocks. Tissue sections of 20 um in thickness were blocked with 5% donkey serum (Jackson ImmunoResearch, 017-000-121) and 0.3% Triton X-100 in PBS (PBST) for 1 hour at room temperature (RT) and then incubated at 4°C overnight with primary antibodies diluted in PBST. The following day, the sections were washed for at least 30 minutes in a Coplin jar of PBS at RT. After the first wash, the sections were incubated with 4′,6-diamidino-2-phenylindole (DAPI, Invitrogen, D1306, 1:1000) and secondary antibodies (Jackson ImmunoResearch, 1:1000) diluted in PBST for an hour at RT. The sections were washed again as previously described. To quench native fluorescence, sections were post-fixed with 4% PFA in PBS for 30 minutes at RT and bleached with a 25% hydrogen peroxide in PBS solution for 20 minutes at RT, after which they were washed in PBS. The sections were mounted with coverslips of #1 thickness (Thermo Scientific, 12541042) and Aqua-Poly/Mount (Polysciences, 18606).

### Wholemount immunostaining

Wholemount immunostaining was performed as previously described.^1,3^ Strips of approximately 3 mm in thickness were cut from the cranial lobe, blocked with 5% donkey serum in PBST for 1 hour on the rocker at RT, and then rocked at 4°C overnight with primary antibodies diluted in PBST. The next day, the strips were washed on the rocker at RT with 1% Tween-20 + 1% Triton X-100 in PBS (PBSTT) for 3 hours, changing the PBSTT every hour. Then, the strips rocked overnight at 4°C with DAPI and secondary antibodies in PBST. On the third day, the strips were washed as previously described and post-fixed in 2% PFA in PBS for at least 2 hours on a RT rocker. The protocol was extended a day for tissue that required native fluorescence to be quenched. After fixation, strips were dehydrated with a methanol/PBS gradient, beginning with 5 minutes in a 50% methanol in PBS solution and then 12 minutes in 100% methanol, changing the methanol every 3 minutes. The strips were then incubated overnight at 4°C in PBS containing 25% hydrogen peroxide. The next day, the bleached strips were rehydrated in a reverse of the previous day’s methanol/PBS gradient and then washed in PBS for at least 10 minutes. Finally, the strips were mounted flatside up with #1 thickness coverslips (Epredia, 12-541-015) and Aqua-Poly/Mount (Polysciences, 18606).

### Confocal imaging

A Nikon Ti2 with AX Confocal System microscope was used for immunofluorescent images. For section immunostaining, Z-stack images were taken at 10 um in thickness while for wholemount, images were taken at 20 um in thickness. For both, step size was set at 0.5 um and imaged using a 20X air objective or 60X oil objective at a resolution of 1024×1024 or 2048×2048. For stitched images of wholemount immunostained strips, images were taken using a 60X oil objective at a resolution of 1024×1024 with pixel dimension 2970 × 1997 × 41.

### Quantification and analysis

ICAM2, CAR4, and PLVAP vasculature surface area was quantified from vessel surface renders using Imaris software. Renderings of ICAM2 and CAR4 vasculature were generated from images of wholemount immunostained strips obtained as described above. PLVAP vasculature, ERG+ nuclei, FOXF1+ nuclei, and Ki67+ nuclei were rendered from images of immunostained sections obtained as described above. At least 3 Z-stacks images per sample were analyzed with Imaris software.

The methodology for quantifying of CAR4 vessel surface area was modified for adult samples to account for the increase in CAR4+ macrophages and the loss of vascular CAR4. Imaris software’s machine learning segmentation (v.10.0.1) was first trained to identify macrophages to render a macrophage-specific surface based on morphology. Then, referencing the original CAR4 channel and the macrophage surface, a new masked channel was created in which voxels within the macrophage surface were assigned a 0 value. A more accurate vessel surface was then rendered from the new channel and quantified.

Quantification of the distal development of CAP2 (CAR4+) vessels in neonate samples was done from stitched images obtained as described above. The maximum intensity projection (MIP) of the images was quantified in Photoshop. The distal edge reference was established using DAPI staining and gridlines every 50 um were used to distribute measurements across the edge. The Photoshop ruler tool was then used to measure, parallel to the vertical gridlines, from the distal edge to the nearest CAR4+ vessel. At least 6 measurements per image and 2-3 images per mouse were measured.

Following quantifications, GraphPad Prism 10 was used to generate plots and perform statistical analysis with Student’s *t*-test. All raw quantifications have been provided (**Dataset S6**).

### Cell dissociation and FACS

Lung dissociation was performed as previously described.^3^ Whole lungs were harvested into Leibovitz’s L-15 Medium (Gibco, 21083027), extra-pulmonary tissues removed, and then minced with forceps. Digestion proceeded in Leibovitz’s with 2 mg/mL of collagenase type I (Worthington, CLS-1 LS004197), 2 mg/mL of elastase (Worthington, ESL LS002294), and 0.5 mg/mL of DNase I (Worthington, D LS002007) at 37°C. After 15 minutes of digestion, the tissue was pipetted to mechanically agitate, then enzymatic digestion continued at 37°C for another 15 minutes. To stop digestion, fetal bovine serum (FBS, Invitrogen, 10082-139) was added to a final concentration of 20%, and the samples were immediately transferred, on ice, to the 4°C cold room. In the cold room, the samples were filtered with a 70 um cell strainer (Corning, 352350) and spun down at 5000 rpm for 1 minute. The resulting supernatant was removed, the cells resuspended in 1 mL of red blood cell lysis (Thermofisher, J62150.AP) for 3 minutes, and then spun down again at 5000 rpm for 1 minute. The red blood cell lysis step was repeated and then the cells were washed with Leibovitz’s with 10% FBS. The samples were then filtered into a 5 mL glass tube with a cell strainer cap (Corning, 352235). The cells were incubated for 30 minutes, on ice, with ECAD-A488 (eBioscience, 53-3249-80, 1:500), ICAM2-A647 (BioLegend, 103114, 1:500), and CD45-PE/Cy7 (Invitrogen, A15452, 1:250). After incubation, the cells were again spun down at 5000 rpm for 1 minute, washed for 5 minutes with Leibovitz’s with 10% FBS, and filtered into a 5 mL glass tube with a cell strainer cap. Finally, the cells were incubated for at least 5 minutes with SYTOX blue (Invitrogen, S34857). The cells were sorted on a BD FACSAria Cell Sorter. After excluding doublets and dead cells, 4 populations were collected: CD45+ cells as the immune lineage, CD45-ICAM2+ as the endothelial lineage, CD45- ICAM2- ECAD+ as the epithelial lineage, and triple negative cells as the mesenchymal lineage.

### Single-cell sequencing and analysis

FACS-purified cells from each lineage were pooled into a single tube per condition (control vs mutant), thereby enriching for the endothelial and mesenchymal lineages. The raw FASTQ data were processed using 10X Genomics CellRanger. Further analysis was performed using R packages Seurat (v.5.4.0) and CellChat (v.2.2.0.9001).^35–37^ Cells were filtered by gene count to exclude those with counts less than 200 or higher than 6000. The mitochondrial gene percentage was also used to filter out cells with more than 10% mitochondrial gene content. Control and mutant data sets were integrated using Seurat’s FindIntegrationAnchors and IntegrateData. The data were log-normalized, variable gene identification performed, scaled, PCA-reduced, and K-means clustered. Cell lineage was established using expression of *Cdh1* (epithelium), *Cdh5* (endothelium), *Ptprc* (immune), and *Col3a1* (mesenchyme). The endothelium was then subsetted and reclustered for analysis of endothelial subpopulations; clusters with nFeature_RNA <2000 were excluded, as well as cluster co-expressing other lineage markers which were identified as doublets. EnhancedVolcano 1.14.0 was used for plotting with a fold change cutoff of 1 and a P value cutoff at 10e-6. CellChat objects were created per sample and merged to allow for cross-sample differential expression analysis. Gene scores were computed from control samples only, and gene lists were generated with the *FindAllMarkers* function after identifying and annotating each cell type. We then selected the top 50 genes per population with an adjusted P-value < 0.05 and a percent expressed of at least 0.5, ranked from highest to lowest logFC. We then use *AddModuleScore* in Seurat to calculate the average expression. The library *enrichR* was used for GO analysis. Seurat was also used to analyze the publicly available human snRNA-seq data from LungMap (LMEX0000004408),^32^ and analyzed as described before.

For scATAC-seq data, Signac (v.1.16.0) and Seurat were used for the analysis.^35,36,38^ TFBSTools (v.1.50.0) and JASPER2020 (v.0.99.10) were used for motif analysis. The whole mouse genome (mm10) was used for gene annotation. Cells were filtered using nucleosome signal, transcriptional start site enrichment score, and peak reads to remove overly noisy and low-quality cells. The data was then run through Signac’s latent semantic indexing (LSI) before LSI dimension reduction and K-means clustering. Cell lineage was established using expression of the markers stated above. The endothelium was then subsetted, and LSI and clustering were rerun for analysis of endothelial subpopulations; clusters co-expressing other lineage markers, which were identified as doublets, were excluded.

### Trajectory analysis

To infer differentiation trajectories among endothelial cell populations in the developing mouse lung, scRNA-seq data were analyzed using CellRank2.^39^ Proliferating ECs were excluded prior to analysis. A k-nearest-neighbor graph and diffusion map were computed in Scanpy, and developmental potential was estimated using the CytoTRACEKernel, which scores cells based on the number of expressed genes as a proxy for stemness. The most stem-like CAP1 cell was designated as the root for pseudotime computation using Palantir (500 waypoints). A PseudotimeKernel was constructed from the resulting pseudotime and used to compute a directed cell–cell transition matrix, which was projected onto the UMAP embedding to visualize directional dynamics across eight annotated endothelial subpopulations. Macrostates were identified using the GPCCA estimator with n = 7 states selected by spectral gap analysis, terminal states were defined from these macrostates, and fate probabilities were computed for each terminal state.

## DATA AND CODE AVAILABILITY

No custom code was generated for this project, but scripts can be shared upon request. Raw sequencing data generated for this manuscript is deposited in GEO; accession number GSE316511. Human snRNA-seq data is available in LungMap (LMEX0000004408).^32^ For time-course mouse scRNA-seq, data from GSE124323, and GSE158205 were used.^1,5^ scATAC-seq data analyzed is available in GSE264098.^20^

## AUTHOR CONTRIBUTIONS

YL and LVE designed the research. YL, AC, DA and JDB performed the research. LVE, YL and LMN performed computational analysis. VVK and TVK provided mice/reagents. YL, AC, LVE and LMN wrote the paper. All authors read and approved the paper.

## ACKNOWLEDGEMENTS

We thank Drs. Vladimir V. Kalinichenko and Tanya V. Kalin (University of Arizona College of Medicine – Phoenix, USA) and Dr. Ralf Adams (University of Münster, Germany) for providing the *Foxf1^CKO^* mice and *Cdh5-CreER*, respectively. This work was supported by the Northwestern University NUSeq Core Facility and the equipment grant S10OD025120. We thank the Robert H. Lurie Comprehensive Cancer Center of Northwestern University in Chicago, IL, for the use of the Flow Cytometry Core Facility, which provided FACS service. The Lurie Cancer Center is supported in part by an NCI Cancer Center Support Grant #P30 CA060553. This work was supported by Northwestern University Start-up Funds and NHLBI R00HL155845 (LVE); K08HL159356 (LMN).

## REFERENCES

1 Vila Ellis, L., et al. Epithelial Vegfa Specifies a Distinct Endothelial Population in the Mouse Lung. Dev Cell 52, 617–630 e616 (2020). 10.1016/j.devcel.2020.01.009

2 Gillich, A. et al. Capillary cell-type specialization in the alveolus. Nature 586, 785–789 (2020). 10.1038/s41586-020-2822-7

3 Vila Ellis, L., et al. p53 maintains lineage fidelity during lung capillary injury-repair in neonatal hyperoxia. JCI Insight 10 (2025). 10.1172/jci.insight.182880

4 Little, D. R. et al. Transcriptional control of lung alveolar type 1 cell development and maintenance by NK homeobox 2-1. Proc Natl Acad Sci U S A 116, 20545–20555 (2019). 10.1073/pnas.1906663116

5 Little, D. R. et al. Differential chromatin binding of the lung lineage transcription factor NKX2-1 resolves opposing murine alveolar cell fates in vivo. Nat Commun 12, 2509 (2021). 10.1038/s41467-021-22817-6

6 Trimm, E. & Red-Horse, K. Vascular endothelial cell development and diversity. Nat Rev Cardiol 20, 197–210 (2023). 10.1038/s41569-022-00770-1

7 Sharma, A. & Niethamer, T. K. Specialized Pulmonary Vascular Cells in Development and Disease. Annu Rev Physiol 87, 229–255 (2025). 10.1146/annurev-physiol-022724-105226

8 Kalucka, J. et al. Single-Cell Transcriptome Atlas of Murine Endothelial Cells. Cell 180, 764–779 e720 (2020). 10.1016/j.cell.2020.01.015

9 Golson, M. L. & Kaestner, K. H. Fox transcription factors: from development to disease. Development 143, 4558–4570 (2016). 10.1242/dev.112672

10 Carlsson, P. & Mahlapuu, M. Forkhead transcription factors: key players in development and metabolism. Dev Biol 250, 1–23 (2002). 10.1006/dbio.2002.0780

11 Mahlapuu, M., Ormestad, M., Enerback, S. & Carlsson, P. The forkhead transcription factor Foxf1 is required for differentiation of extra-embryonic and lateral plate mesoderm. Development 128, 155–166 (2001). 10.1242/dev.128.2.155

12 Dharmadhikari, A. V. et al. Lethal lung hypoplasia and vascular defects in mice with conditional Foxf1 overexpression. Biol Open 5, 1595–1606 (2016). 10.1242/bio.019208

13 Stankiewicz, P. et al. Genomic and genic deletions of the FOX gene cluster on 16q24.1 and inactivating mutations of FOXF1 cause alveolar capillary dysplasia and other malformations. Am J Hum Genet 84, 780–791 (2009). 10.1016/j.ajhg.2009.05.005

14 Towe, C. T. et al. Infants with Atypical Presentations of Alveolar Capillary Dysplasia with Misalignment of the Pulmonary Veins Who Underwent Bilateral Lung Transplantation. J Pediatr 194, 158–164 e151 (2018). 10.1016/j.jpeds.2017.10.026

15 Yost, C. E. et al. A Long-Term Survivor With Alveolar Capillary Dysplasia. JACC Case Rep 2, 1492–1495 (2020). 10.1016/j.jaccas.2020.05.055

16 Sirianansopa, K., Prasertsan, P., Ruangnapa, K., Saelim, K. & Kor-Anantakul, P. Unusual presentation of alveolar capillary dysplasia with misalignment of the pulmonary veins in a child with respiratory syncytial virus pneumonia: A case report. Respirol Case Rep 11, e01089 (2023). 10.1002/rcr2.1089

17 Pradhan, A. et al. The S52F FOXF1 Mutation Inhibits STAT3 Signaling and Causes Alveolar Capillary Dysplasia. Am J Respir Crit Care Med 200, 1045–1056 (2019). 10.1164/rccm.201810-1897OC

18 Wang, G. et al. Endothelial progenitor cells stimulate neonatal lung angiogenesis through FOXF1-mediated activation of BMP9/ACVRL1 signaling. Nat Commun 13, 2080 (2022). 10.1038/s41467-022-29746-y

19 Kalinichenko, V. V. et al. Defects in pulmonary vasculature and perinatal lung hemorrhage in mice heterozygous null for the Forkhead Box f1 transcription factor. Dev Biol 235, 489–506 (2001). 10.1006/dbio.2001.0322

20 Hassan, D. & Chen, J. CEBPA restricts alveolar type 2 cell plasticity during development and injury-repair. Nat Commun 15, 4148 (2024). 10.1038/s41467-024-48632-3

21 Ren, X. et al. FOXF1 transcription factor is required for formation of embryonic vasculature by regulating VEGF signaling in endothelial cells. Circ Res 115, 709–720 (2014). 10.1161/CIRCRESAHA.115.304382

22 Ren, X. et al. Postnatal Alveologenesis Depends on FOXF1 Signaling in c-KIT(+) Endothelial Progenitor Cells. Am J Respir Crit Care Med 200, 1164–1176 (2019). 10.1164/rccm.201812-2312OC

23 Narvaez Del Pilar, O., Gacha Garay, M. J. & Chen, J. Three-axis classification of mouse lung mesenchymal cells reveals two populations of myofibroblasts. Development 149 (2022). 10.1242/dev.200081

24 Hoggatt, A. M. et al. The transcription factor Foxf1 binds to serum response factor and myocardin to regulate gene transcription in visceral smooth muscle cells. J Biol Chem 288, 28477–28487 (2013). 10.1074/jbc.M113.478974

25 Wang, Y. et al. Ephrin-B2 controls VEGF-induced angiogenesis and lymphangiogenesis. Nature 465, 483–486 (2010). 10.1038/nature09002

26 Saul, D. et al. A new gene set identifies senescent cells and predicts senescence-associated pathways across tissues. Nat Commun 13, 4827 (2022). 10.1038/s41467-022-32552-1

27 Freund, A., Laberge, R. M., Demaria, M. & Campisi, J. Lamin B1 loss is a senescence-associated biomarker. Mol Biol Cell 23, 2066–2075 (2012). 10.1091/mbc.E11-10-0884

28 Tian, W. et al. An embryonic artery-forming niche reactivates in pulmonary arterial hypertension. bioRxiv, 2025.2005.2002.651303 (2025). 10.1101/2025.05.02.651303

29 Nguyen, V. D. & Zhou, B. From Heterogeneity to Plasticity: Endothelial Dynamics in Lung Disease. Pulmonary Circulation 16, e70287 (2026). 10.1002/pul2.70287

30 Wang, H. et al. Multiomic profiling reveals early disease mechanisms of lung fibrosis progression. ERJ Open Research 12, YI501 10.1183/23120541.Lsc-2026.Yi501

31 Shirazi, S. P. et al. Bronchopulmonary dysplasia with pulmonary hypertension associates with semaphorin signaling loss and functionally decreased FOXF1 expression. Nat Commun 16, 5004 (2025). 10.1038/s41467-025-60371-7

32 Guo, M. et al. Single Cell Multiomics Identifies Cells and Genetic Networks Underlying Alveolar Capillary Dysplasia. Am J Respir Crit Care Med 208, 709–725 (2023). 10.1164/rccm.202210-2015OC

33 Sen, P., Thakur, N., Stockton, D. W., Langston, C. & Bejjani, B. A. Expanding the phenotype of alveolar capillary dysplasia (ACD). J Pediatr 145, 646–651 (2004). 10.1016/j.jpeds.2004.06.081

34 Mo, A. et al. Epigenomic Signatures of Neuronal Diversity in the Mammalian Brain. Neuron 86, 1369–1384 (2015). 10.1016/j.neuron.2015.05.018

35 Hao, Y. et al. Dictionary learning for integrative, multimodal and scalable single-cell analysis. Nat Biotechnol 42, 293–304 (2024). 10.1038/s41587-023-01767-y

36 Zhang, L. et al. Single-cell transcriptomic profiling of lung endothelial cells identifies dynamic inflammatory and regenerative subpopulations. JCI Insight 7 (2022). 10.1172/jci.insight.158079

37 Jin, S. et al. Inference and analysis of cell-cell communication using CellChat. Nat Commun 12, 1088 (2021). 10.1038/s41467-021-21246-9

38 Stuart, T., Srivastava, A., Madad, S., Lareau, C. A. & Satija, R. Single-cell chromatin state analysis with Signac. Nat Methods 18, 1333–1341 (2021). 10.1038/s41592-021-01282-5

39 Weiler, P., Lange, M., Klein, M., Pe’er, D. & Theis, F. CellRank 2: unified fate mapping in multiview single-cell data. Nat Methods 21, 1196–1205 (2024). 10.1038/s41592-024-02303-9

