## Supplemental Figures for "*Foxf1* is required for the specification and maintenance of pulmonary capillary identity"

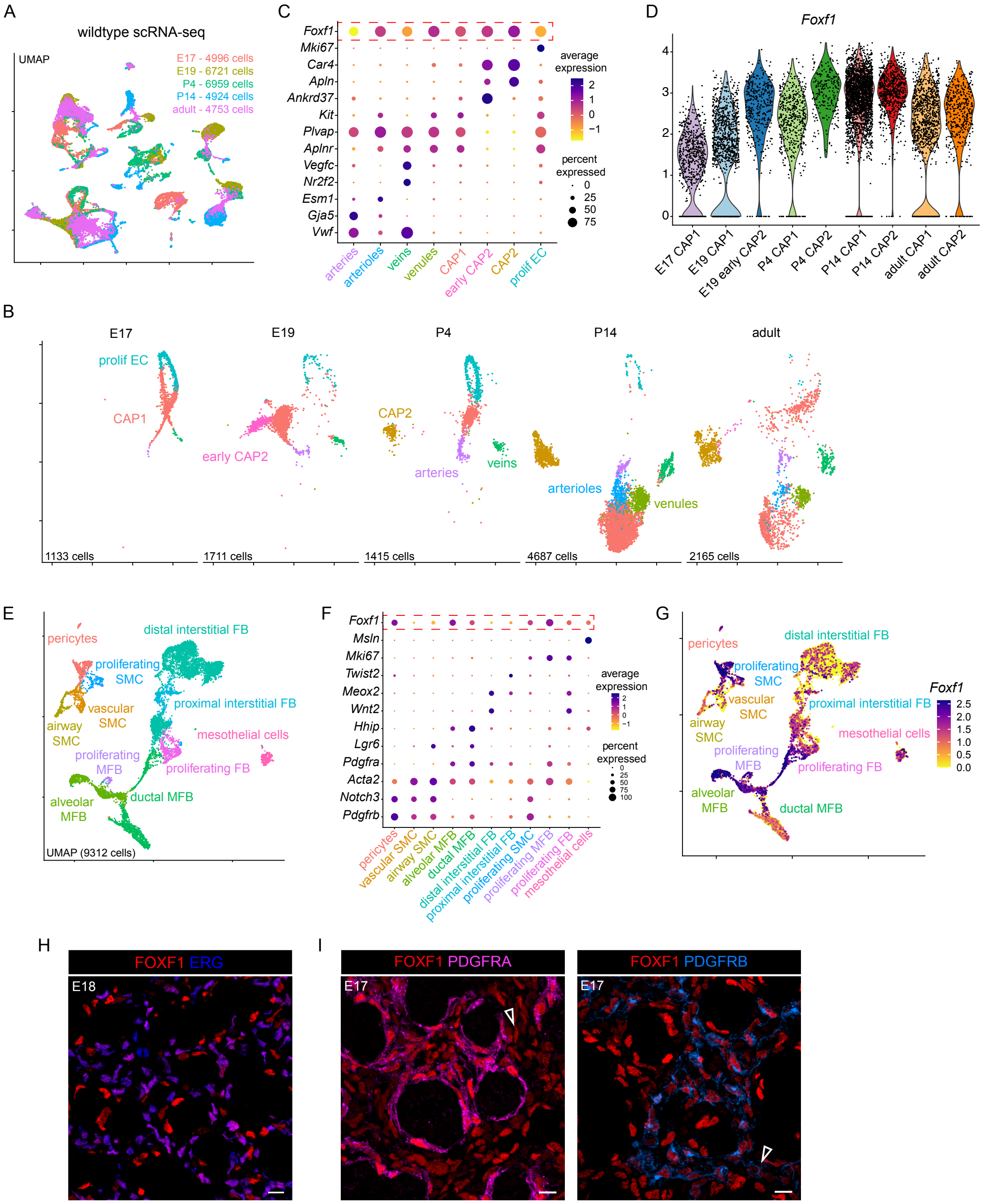


**Fig. S1.** (**A**) UMAP of time-course scRNA-seq data from wildtype lungs, colored by time point. (**B**) UMAPs of endothelial-lineage cells of wildtype time-course scRNA-seq data, colored by subpopulation. (**C**) Dot plot of markers used to identify endothelial subpopulations; *Foxf1* expression is highlighted by a red dash box. (**D**) Violin plot showing how the level of *Foxf1* expression changes in capillary ECs across developmental time. (**E**) UMAP of mesenchymal cells of wildtype time-course scRNA-seq data, colored by subpopulation. (**F**) Dot plot of markers used to identify mesenchymal subpopulations; *Foxf1* expression is highlighted in red dash box. (**G**) Feature plot of *Foxf1* expression in mesenchymal cells showing high expression in pericytes and myofibroblasts. (**H**) Section immunostaining of wildtype E18 lung showing endothelial (ERG+) FOXF1 expression. (**I**) Section immunostainings of wildtype E17 lung showing FOXF1, PDGFRA (fibroblasts, open arrowhead) or PDGFRB (pericytes, open arrowhead). E, embryonic day; prolif, proliferative; FB, fibroblasts; MFB, myofibroblasts; SMC, smooth muscle cells. Scale bars = 10 μm.

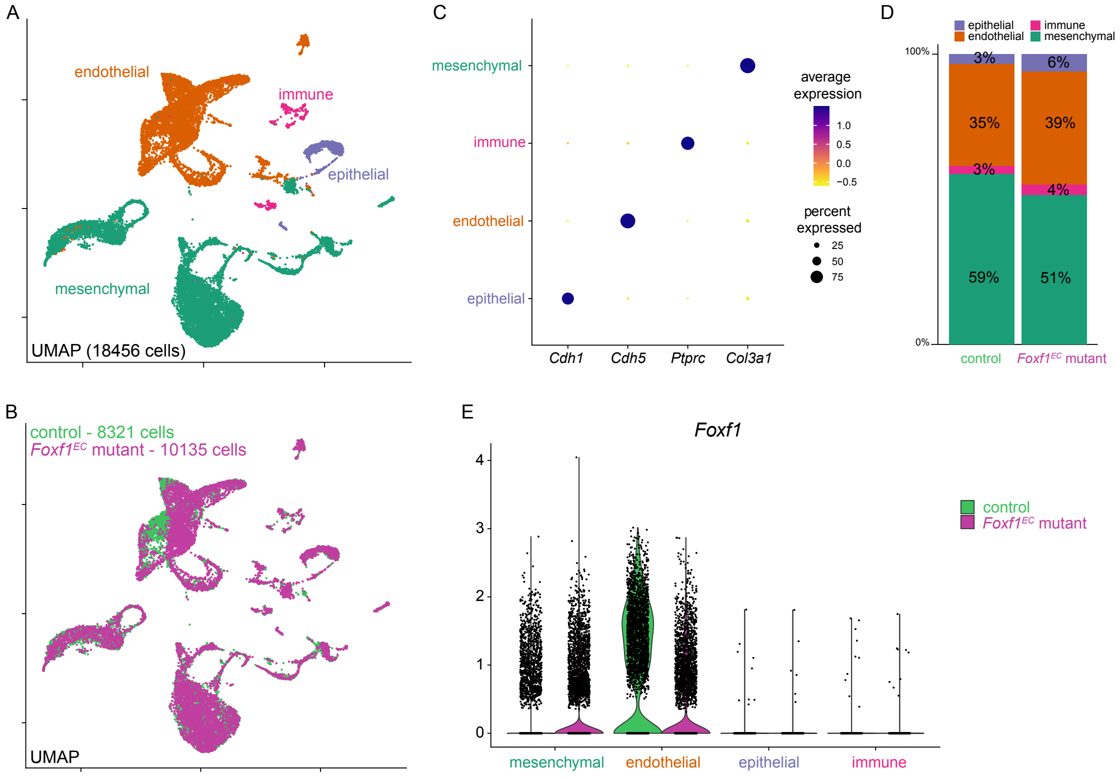


**Fig. S2. (A)** UMAP showing all cells in our scRNA-seq data between P4 control and *Foxf1*$^{EC}$ mutants, colored by lineage or **(B)** colored by condition. **(C)** Dot plot of markers used to identify lineages. **(D)** Stacked barplot showing percentages of lineage populations between control and *Foxf1*$^{EC}$ mutant. **(E)** Violin plot confirming decrease in endothelial *Foxf1* transcripts in *Foxf1*$^{EC}$ mutant.

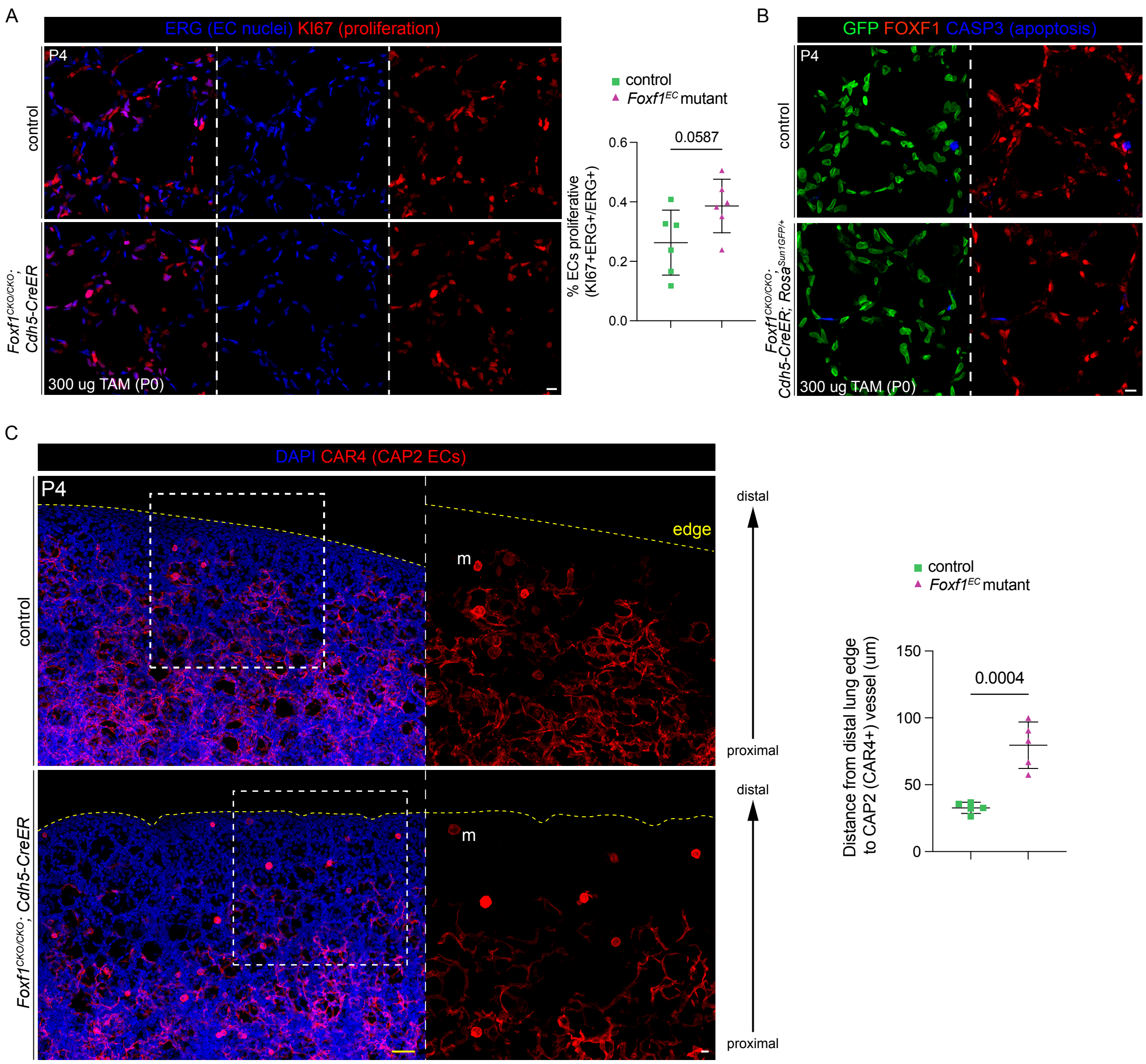


**Fig. S3. (A)** Section immunostaining of P4 lungs showing proliferation marker KI67 and ERG. Quantification of the percent of ECs that are proliferative show no significant increase in the *Foxf1*$^{EC}$ mutant. **(B)** Section immunostaining of P4 lungs showing no increase in cell death (cleaved-CASP3) overall or in ECs (*Rosa*$^{SUN1GFP}$ labels ECs, GFP+ nuclei). **(C)** Wholemount immunostaining of P4 lungs showing DAPI and CAR4 in the distal region of the lung. The *Foxf1*$^{EC}$ mutant has significantly reduced CAR4 vessel development towards the distal edge (dashed yellow line) as quantified in the plot. Each quantification datapoint represents the average of 3 distinct images within 1 mouse. All *p*-values calculated using Student's t test. Boxed regions are magnified. TAM, tamoxifen; m, macrophage; P, postnatal day. Scale bars in A, B, and inset of C = 10 μm; scale bar in C = 50 μm.

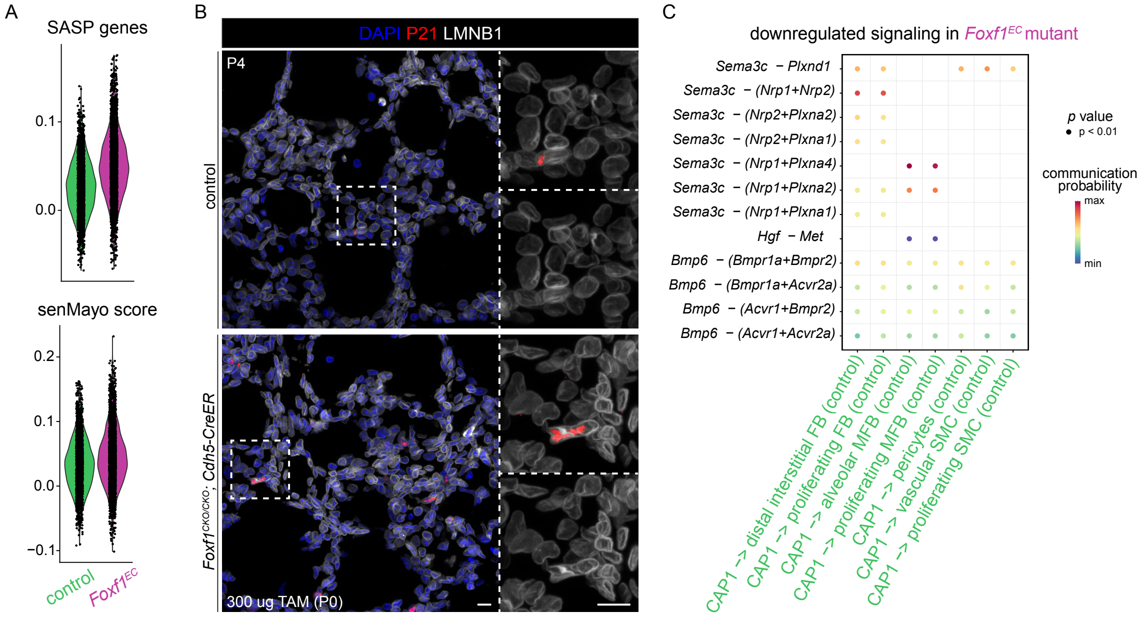


**Fig. S4. (A)** Violin plot of module scores generated from the senescence-associated secretory protein (SASP) gene list and senMayo senescent gene list. **(B)** Section immunostaining of P4 lungs showing no loss of LMNB1 in cells expressing P21. **(C)** Dot plot of predicted, downregulated ligand-receptor interactions in *Foxf1*$^{EC}$ mutant with CAP1 as sender and mesenchymal subpopulations as targets. Loss of predicted interactions should be interpreted cautiously, as severe downregulation of CAP1-associated genes may reduce ligand or receptor expression below the threshold required for interaction inference. Boxed regions are magnified. TAM, tamoxifen; P, postnatal day; mut, mutant; prolif, proliferative; FB, fibroblasts; MFB, myofibroblasts; SMC, smooth muscle cells. Scale bars = 10 μm.
